# HDAC inhibition unlocks tumor plasticity and enhances immunotherapy response in Myc-Driven Small Cell Lung Cancer

**DOI:** 10.1101/2025.08.06.668958

**Authors:** Azam Ghafoor, Linying Zhu, Zoe Weaver Ohler, Rajaa El Meskini, Devon Atkinson, Amanda Day, Laura Bassel, Weixin Wang, Haoxuan Ying, Katherine R. Calvo, Lorinc Pongor, Yves Pommier, Anish Thomas, Yilun Sun

## Abstract

Small Cell Lung Cancer (SCLC) is a highly aggressive malignancy, accounting for approximately 15% of all lung cancer cases. Characterized by low immunogenicity, SCLC may utilize epigenetic mechanisms to evade immune detection. Here, we demonstrate that entinostat, a class I histone deacetylase inhibitor (HDACi) upregulates immune-related genes in human SCLC cells. In vivo, we confirmed entinostat treatment increased expression of immunecheckpoint ligands and antigen presentation machinery in Myc-driven tumors in a Rb1/Trp53/Myc^T58A^ (RPM) SCLC mouse model, while shifting tumors from a neuroendocrine(NE)-high to a NE-low phenotype. Notably, combining entinostat with anti-PD-1 immunotherapy significantly enhances T-cell infiltration, suppresses tumor growth, and prolongs survival in RPM allograft models. These findings underscore the potential of entinostat to reprogram the immunological landscape and NE status of SCLC, enhance immune checkpoint blockade efficacy, and improve therapeutic outcomes.

## INTRODUCTION

Small cell lung cancer (SCLC) is an aggressive, high-grade neuroendocrine carcinoma with a dismal prognosis, representing approximately 15% of all lung cancer diagnoses.^1–3^ Most SCLCs exhibit classic neuroendocrine (NE) features with positive NE markers such as synaptophysin, chromogranin A, and CD56; while a smaller subset presents as non-neuroendocrine variants.

Based on transcriptional profiling, SCLC can be further categorized into four molecular subtypes: SCLC-A (ASCL1), SCLC-N (NEUROD1) SCLC-P (POU2F3) and SCLC-I (lack expression in A/N/P). SCLC-A and-N typically exhibit NE-high phenotypes, while SCLC-P and-I correspond to NE-low states.^4^

Despite initial chemosensitivity, SCLC patients frequently relapse with refractory disease. Standard treatment regimens, unchanged for decades, primarily involve DNA-damaging chemotherapeutics.^5^ Recently, the incorporation of immune checkpoint inhibitors, such as anti-PDL1 monoclonal antibodies atezolizumab or durvalumab, into the first-line treatment combining platinum-based chemotherapy with topoisomerase II inhibitor etoposide, has yielded modest improvements in survival. Although initial response rates are remarkably high (60–70%), the majority of patients relapse within one year. The median progression-free survival (PFS) in contemporary clinical trials remaining around five months.^6,7^ Second-line treatments for extensive-stage SCLC include topoisomerase I inhibitor topotecan and transcription inhibitor lurbinectedin. While lurbinectedin has demonstrated a higher overall response rate of 35% compared to topotecan (25%),^8^ both agents offer limited clinical benefit. Furthermore, owing in part due to the low immunogenicity of SCLC tumors, immune checkpoint monotherapies have shown minimal efficacy, with consistently low response rates across multiple clinical studies.^9^

The FDA approved CD3-DLL3 T-cell engager tarlatamab directs autologous T-cells to cancer cells expressing delta-like ligand for patients with SCLC in the third-line setting. The response rate in phase 2-3 trials was only 35-40%, and the median PFS remained at less than 5 months^10,11^ The low response rate, with only 1% achieving a complete response, is partly attributed to the expression of inhibitory immune modulators, a transcriptional shift toward a less neuroendocrine (NE) subtype, characterized by reduced levels of ASCL1 or NEUROD1, and the downregulation of antigen processing and presentation pathways.^12–14^ These changes collectively suppress T cell activity and contribute to resistance to tarlatamab.NE-high subtypes (A and N) typically respond well to platinum-based chemotherapy but frequently relapse with chemo-resistant phenotypes.

Conversely, NE-low subtypes (P and Y) exhibit greater immune infiltration, such as elevated CD8+ T cells and higher PD-L1, indicating potential susceptibility to immune checkpoint inhibitors.^15^ Intramural heterogeneity and subtype switching under treatment pressure present therapeutic challenges. These challenges underscore an urgent need for novel combination-based epigenetic approaches to drive tumor plasticity into a more favorable NE low phenotype and enhance immunotherapy markers to improve outcomes in refractory SCLC.^16^

Despite harboring highly abnormal genomes with extensive mutational burdens, SCLC tumors are non-immunogenic and typically exhibit low expression of PD-L1 and the antigen-presenting MHC class I.^17^ Emerging evidence suggests that post-translational histone modifications, histone deacetylation in particular, play a crucial role in suppressing immune-related gene expression, including PD-L1 and antigen-presenting molecules.^18^ Preclinical studies have shown that histone deacetylase (HDAC) inhibition synergizes with PD-1 blockade and hence improves its antitumor activity in multiple murine tumors.^19–21^ HDAC inhibition appears to modulate myeloid-derived suppressor cells (MDSCs) by reducing their immunosuppressive activity and fostering a more favorable immune milieu.^20^

In this study, we investigate the role of the Class I HDAC inhibitor entinostat as a means of epigenetically reprogramming SCLC to an immunogenic state. We show that entinostat upregulates immune-related genes and antigen presentation machinery in both in vitro and in vivo SCLC models. Additionally, entinostat suppresses neuroendocrine lineage drivers, enhances the infiltration of immune cells, and improves the efficacy of anti-PD-1 immunotherapy. The combination of entinostat and anti-PD-1 significantly suppresses tumor growth and prolongs survival in RPM allograft models. These findings, taken together, support the rationale for combining epigenetic therapy with immune checkpoint blockade in Myc driven SCLC, which occur in 20% of SCLC and confer a more aggressive phenotype and poor prognosis.^22,23^

## RESULTS

### Immune checkpoint and MHC genes are repressed in small cell lung cancers

While cancer immunotherapy targeting immune checkpoints such as PD-1/PD-L1 yielded clinical benefits in a spectrum of malignancies, its benefit in SCLC remains limited. Several barriers may contribute to the low efficacy of immune checkpoint inhibitors in SCLC: (1) substantial heterogeneity within the tumor and microenvironment;^24^ (2) significantly reduced PD-L1 expression in SCLC compared to lung adenocarcinoma and other solid tumors (Fig. 1A). This low baseline expression indicates an inactive immune microenvironment and reduced likelihood to respond to immune checkpoint inhibitors; and (3) broad downregulation of MHC class I genes (HLA-A, HLA-B, HLA-C) which present endogenous antigens for immune surveillance, and the antigen-processing gene TAP1 (Fig. 1B, Fig. S1A) which are essential for antigen presentation and immune recognition. The repression of MHC-I and antigen-processing machinery genes is a well-documented mechanism of both intrinsic and acquired resistance to immunotherapy in various cancers.^25–27^ Therefore, increasing PD-L1, restoring MHC-I expression and antigen presentation may enhance immune recognition by T cells and improve the efficacy of immunotherapy in SCLC.

**Figure 1.**
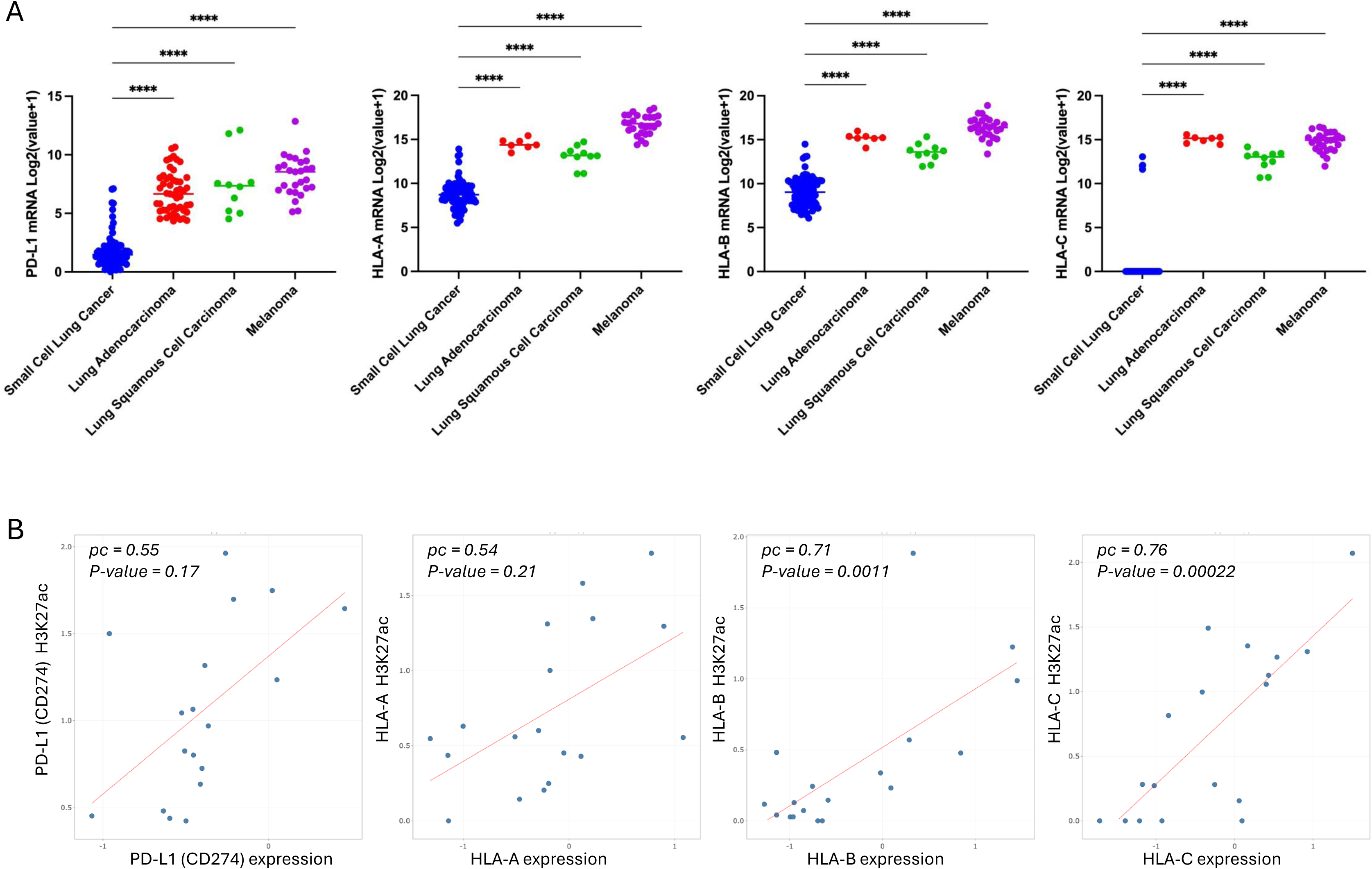
PD-L1 and MHC genes are repressed in SCLC amongst solid malignancies. A. Gene expression levels of *PD-L1, HLA-A, HLA-B* and *HLA-*C in different cancer types. Data is from CBioPortal (https://www.cbioportal.org/). **** p < 0.0001. B. Correlation between H3K27ac ChIP-seq signal and gene expression of *PD-L1 (CD274), HLA-A, HLA-B, HLA-C* in SCLC cell lines. Plot is generated with the sclcCellMinerCDB (https://discover.nci.nih.gov/rsconnect/SclcCellMinerCDB/).

To identify potential transcriptional regulators of PD-L1 and MHC-I expression in SCLC, we examined correlations between PD-L1/MHC gene levels and the levels of epigenetic markers including histone acetylation (H3K27ac) and DNA methylation at promoter regions (Fig. 1C). We observed a strong positive correlation between the gene expressions of PDL1 and HLA markers and histone acetylation (H3K27ac) but not DNA methylation (Fig. S1B). These results suggest that histone acetylation and deacetylation play a crucial role in the epigenetic regulation of immune genes in SCLC, potentially providing a therapeutic avenue to enhance immunogenicity.

### Entinostat upregulates PD-L1 and MHC expression in small cell lung cancer cell lines

Entinostat is a benzamide class I-selective histone deacetylase inhibitor (HDAC 1, 2, 3-nuclear) undergoing clinical trials for the treatment of various cancers.^20,28–31^ To determine if HDAC inhibition by entinsostat improves expression of immune markers in SCLC, we treated a panel of SCLC cell lines, H889, H209, H82, H524, and DMS-114, with entinostat and found that the treatment elevated mRNA levels of PD-L1, MHC-I, and MHC-II in a dose-dependent manner (Fig. 2A and Fig. S2A). Consistently, protein expression levels of PD-L1 and MHC-I are also upregulated in a dose-dependent manner following entinostat treatment (Fig. 2B). Notably, the NE-low cell line DMS-114 showed limited induction of HLA-C and A genes and less dose-dependent changes on HLA-B genes upon entinostat treatment, suggesting that neuroendocrine status may influence entinostat responsiveness. Further, entinostat was found to induce antigen-processing and presentation genes TAP1 and PSMB8 and key anti-tumor immune-stimulatory chemokines CXCL10 and IFNγ in a dose-dependent manner (Fig. S2B and S2C), indicating potential to enhance adaptive immune responses.

**Figure 2.**
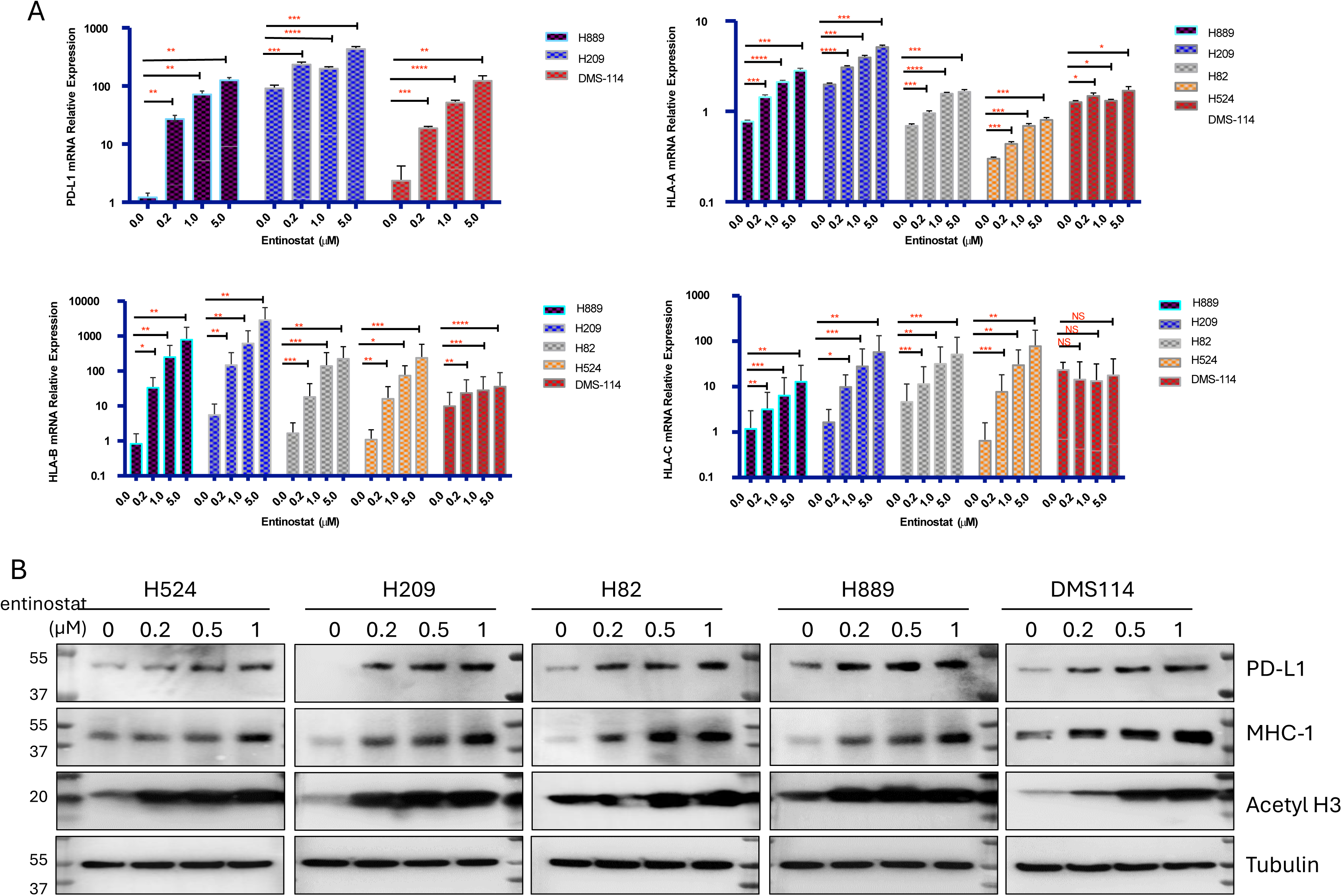
HDAC inhibitor enthinostat regulates PD-L1 and MHC genes expression in SCLC cell lines. A. The relatve gene expression of *D274 (PD-L1), HLA-A, HLA-B* and *HLA-C* on SCLC cell lines H889, H209, H82, H524, and DMS-114 treated with increasing dose (0 uM, 0.2 uM, 1 uM and 5 uM) of entinostat for 24 hours. The relative gene expression was determined by quantitative real-time PCR assay. Error bars are representative of technical triplicates. *p<0.05, **p<0.01, ***p<0.001, ****p<0.0001. B. Western blot to show the protein level of PD-L1, MHC-1, Acetylated histone 3 (target) and tubulin (control) on the above five SCLC cell lines treated with entinostat for 24 hours in increasing dose of 0uM, 0.2uM, 0.5uM and 1uM.

### RPM mouse model recapitulates NE-high, immune-cold small cell line cancers

We next sought to investigate the effect of entinostat on immune gene regulation in an in vivo lung tumor model with low immunogenicity.(Meuwissen et al., 2003; Oser et al., 2024; Rudin et al., 2021) Inactivation of Rb1 and Trp53, combined with the expression of MYC^T58A^, an oncogenic form of MYC, leads to the development of NE-high SCLC.^22,32^ MYC overexpression, often via extrachromosomal amplification, is observed in approximately 20% of SCLC patients and is associated with more aggressive behavior.^33,34^ We used the Rb1/Trp53/Myc^T58A^ (RPM) genetically engineered mouse (GEM) model to recapitulate human SCLC (Fig. 3A),^35^ which developed NEUROD1-high primary lung tumors with rapid onset. Tumor progression in the RPM model is characterized by a steep decline in survival and rapid tumor progression at around 8 weeks (Fig. 3B). The RPM mice exhibited complete opacification of the hemithorax at around 9 weeks as detected by magnetic resonance imaging (MRI) (Fig. S3A). Histopathological analyses confirmed that the induced tumor possesses human SCLC features, including small cell morphology, scant cytoplasm, absent nucleoli, neoplastic cells arranged in sheets and ribbons, and tumors originating in the large bronchi (Fig. S3B). Frequent lobular coagulative necrosis was also observed, indicating tumor-associated vascular occlusion (Fig. S3B-C). The RPM tumors express NEUROD1 and high levels of *CD56*, another neuroendocrine SCLC marker (Fig. 3C).

**Figure 3.**
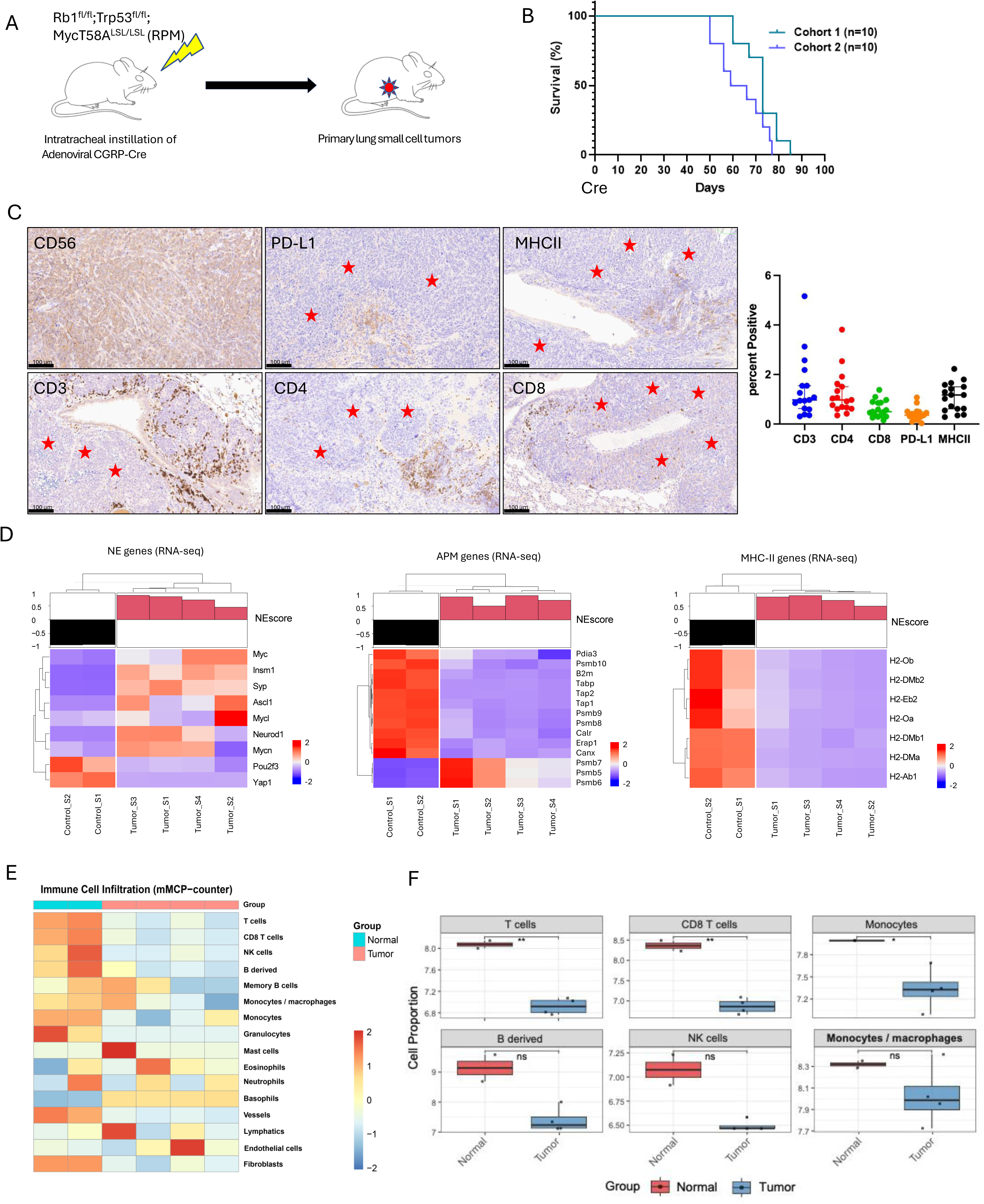
Evaluation of immune cell population in SCLC RPM GEM tumor model. A. *Rb1/Trp53/Myc^T58A^* (RPM) mice weer induced at 8 weeks of age with adenoviral Cgrp-Cre (2.5 × 10^7^ pfu). B. Survival of RPM mice was consistent over 2 separate cohorts. C. Immunohistochemistry staining (left) and automated quantification (right) of *NCAM(CD56), PD-L1, MHC-II, CD3, CD4* and *CD8*. Scale bar is 10 uM (20x). Red star mark tumor regions. D. Gene expression heatmap of lungs from two uninduced mice and 4 RPM tumors on the indicated gene sets. Top of the heatmap is the NE score for each sample. E. Immune cell populations in normal and tumor groups analyzed with mMCP-counter. F. Example of the differential distribution of the indicated immune cell populations between normal and tumor samples from mMCP-counter analysis.

NE-high SCLC exhibits an “immune desert” tumor microenvironment characterized by poor immune cell infiltration, whereas the NE-low subtype often shows enhanced immune cell infiltration.^36^ Immune profiling of RPM tumors through IHC revealed low T-cell infiltration (by staining for T cell markers *CD3, CD4*, and *CD8*) and low PD-L1 and MHC-II expression relative to tumor-adjacent normal tissues, indicative of an inactive immune microenvironment (Fig. 3C).

We next performed RNA sequencing (RNA-seq) of RPM lung tumors compared to normal lung tissues from uninduced mice to further assess the NE characteristics (Fig. 3D, top) While individual tumors displayed heterogeneous NE gene expressions, all the tumors exhibited higher levels of NE markers (*Neurod1, Ascl1, Insm1,* and *Syp*) and lower levels of Yap1 compared to adjacent normal lung tissue (Fig. 3D, left). In addition, RPM tumors exhibited low expression of antigen processing machinery (APM) genes, a determinant of tumor response to immune checkpoint blockade,^37,38^ compared to normal tissues (Fig. 3D, middle). MHC-II genes, which are primarily expressed in antigen-presenting cells (APCs) and important for tumor antigen presentation, were also suppressed compared to normal tissues (Fig. 3D, right). These results suggest that SCLC evades immune evasion through suppressing APM and MHC genes.

To further compare immune cell subsets in the tumor immune microenvironment and normal tissues in RPM models, we used the murine Microenvironment Cell Populations-counter (mMCP-counter), to quantify immune and stromal cell populations in heterogeneous murine samples.^39^ This analysis revealed lower infiltration of total T cells, CD8+ T cells, and monocytes in lung tumors versus normal tissue (Fig. 3E - F). The analysis confirmed that T cells (p<0.01), CD8+ T cells (p<0.01), and monocytes (p<0.05) were significantly higher in normal tissues than in the tumor. NK cells, B-derived populations displayed higher infiltration in normal tissues than in tumor tissues. Collectively, histopathological and transcriptomic analyses confirm that RPM tumors recapitulate classical NE-high phenotype with tumor cells actively suppressing immunogenic markers to evade immune system. This discovery can be used to develop therapeutic strategies for unlocking tumor plasticity in SCLC and drive adaptive immunity to improve anti-tumor immune responses in SCLC.

### Entinostat treatment shifts NE-high tumors toward NE-low and enhances immunogenicity

Prompted by our in-vitro observations that entinostat upregulates immune checkpoint, APM, and antigen presentation genes in SCLC cell lines, we hypothesized that the drug could shift SCLC tumors from a low immunogenic state (NE-high) to a high immunogenic state (NE-low), thereby enhancing their response to immune checkpoint blockade. To test this hypothesis, we treated RPM mice using entinostat alone, anti-PD-1 alone, or a combination of both (Fig. 4A). We initially conducted a time-course experiment mimicking drug dosing and clearance in a SCLC cell line and observed that PD-L1 expression peaks at 24 hours following entinostat treatment and declines rapidly upon drug washout (Fig. S4A). This finding suggests that PD-L1 induction by entinostat is reversible and suggests concurrent daily dosing schema rather than sequential dosing of entinostat with immune checkpoint blockade to maximize therapeutic synergy.

**Figure 4.**
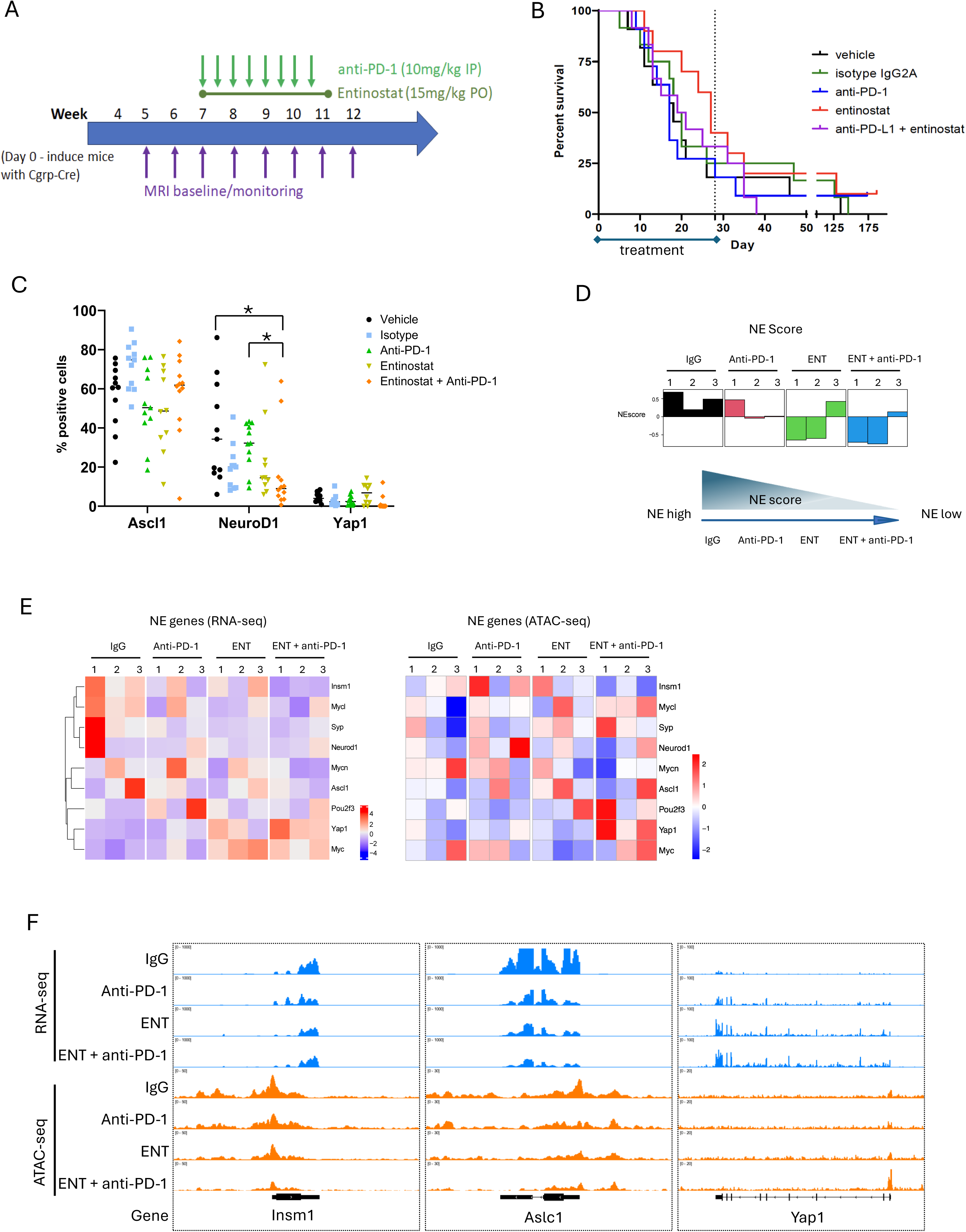
Entinostat change the RPM tumors neuroendocrine status. A. Schematic of the entinostat and anti-PD-1 treatment regimen in RPM mice. PO: Oral gavage. IP: intraperitoneal. MRI: Magnetic resonance imaging. B. Survival curve of the control and treatment groups. P-value is not significant for any treatment compared to vehicle. C. Quantification of IHC staining of Ascl1, Neurod1 and Yap1 in each group of tumors. D. Neuroendocrine score of each sample in four groups. E. Heatmap of NE-associated genes on their expression level and ATAC-seq peak signal in these gene’s promoter region (2kb upstream and downstream of TSS) F. RNA-seq peaks and ATAC-seq peaks track signals on the NE genes.

Evaluation of drug toxicity confirmed that entinostat, both as a monotherapy at 15mg/kg and in combination with 10 mg/ml anti-PD-1, were well-tolerated. No mice met the criteria for euthanasia (>20% body weight loss or signs of toxicity) using entinostat monotherapy and no abnormalities were noted by gross necropsy. Maximum body weight loss for a single mouse was 5% in the anti-PD-1 treated group and 8% in the combination treated group (Fig. S4B). The mice did not exhibit additional signs of toxicity.

To evaluate efficacy of combination treatment in tumor-bearing mice, induced RPM GEM were treated with entinostat 15mg/kg PO once daily for 28 days, and anti-PD-1 was injected intraperitoneally twice per week (days 1 and 4) at 10mg/kg x 4 weeks. Primary SCLC tumors were monitored for growth using MRI starting at week 5 and treatment was administered at week 7. Due to the high inter-sample variability and rapid tumor growth within treatment groups, we did not observe a statistically significant reduction in tumor volume nor improvement in survival rates in the RPM model using the combination therapy when compared to monotherapy or vehicle (Fig. 4B and S4C–D). This was likely due to rapid tumor growth from MYC-driven primary tumors situated in lung and bronchus.

Despite the lack of observable tumor growth inhibition, immunohistochemical analyses showed a reduction of median NEUROD1expression in tumors from mice treated with entinostat monotherapy and in tumors treated with the combination therapy (Fig. 4C). As a validation, we performed RNA-seq on a subset of tumors from each treatment group to assess the NE score for based on the publihsed 50-gene expression signature.^40^ Entinostat, both alone and in combination with anti-PD-1, resulted in a substantial reduction in NE score (Fig. 4D). To gain further mechanistic insight into this NE status transition, we analyzed NE-associated gene expression by RNA-seq and chromatin accessibility by ATAC (Assay for Transposase-Accessible Chromatin)-seq. RNA-seq showed downregulation of NE-high markers *Neurod1, Insm1, Syp, and Ascl1* and upregulation of NE-low markers *Yap1 and Myc* (Fig. 4E, left). ATAC-seq revealed that these gene expression changes were accompanied by chromatin remodeling at the promoter sites, with closure of NE-high gene (e.g., *Insm1*) promoters and opening of NE-low gene loci (e.g., *Yap1*) with combination treatments (Fig. 4E, right, and Fig. 4F). These results suggest that MYC-driven SCLC can transition from a NE-high to NE-low state through HDAC inhibition (HDACi), a process primarily governed by changes in chromatin accessibility at NE-associated genes via histone acetylation. Hence, this important discovery holds therapeutic relevance for targeting lineage plasticity in SCLC, offering a potential strategy to overcome NE resistant immunotherapies and improve patient outcomes.

### Entinostat in combination with anti-PD-1 treatment activates antigen presentation

Next, we assessed the effect of entinostat in combination with anti-PD-1 treatment on immune activation genes. APM genes, including *Tap1, Tap2, B2m, Tapbp, Psmb8, Psmb9, Erap1,* and *Carnx*, which are repressed in RPM tumors, were activated following the combination treatment (Fig. 5A, left). Of note, the proteasome genes *psmb8* and *psmb9,* but not *psmb5, psmb6, psmb7,* and *psmb10* genes, were upregulated following the treatments. *Psmb5, psmb6, and psmb7* genes encode constitutive proteasome components, whereas *psmb8, psmb9, and psmb10* encode immunoproteasome components, which have been associated with increased tumor-infiltrating lymphocytes and longer survival.^41^ The increased expression of genes encoding immunoproteasome suggests higher infiltration and immunogenic status. ATAC-seq showed increased chromatin accessibility at the loci of these APM genes following the treatment of entinostat plus anti-PD-1 (Fig. 5A right, 5C, and Fig. S5A).

**Figure 5.**
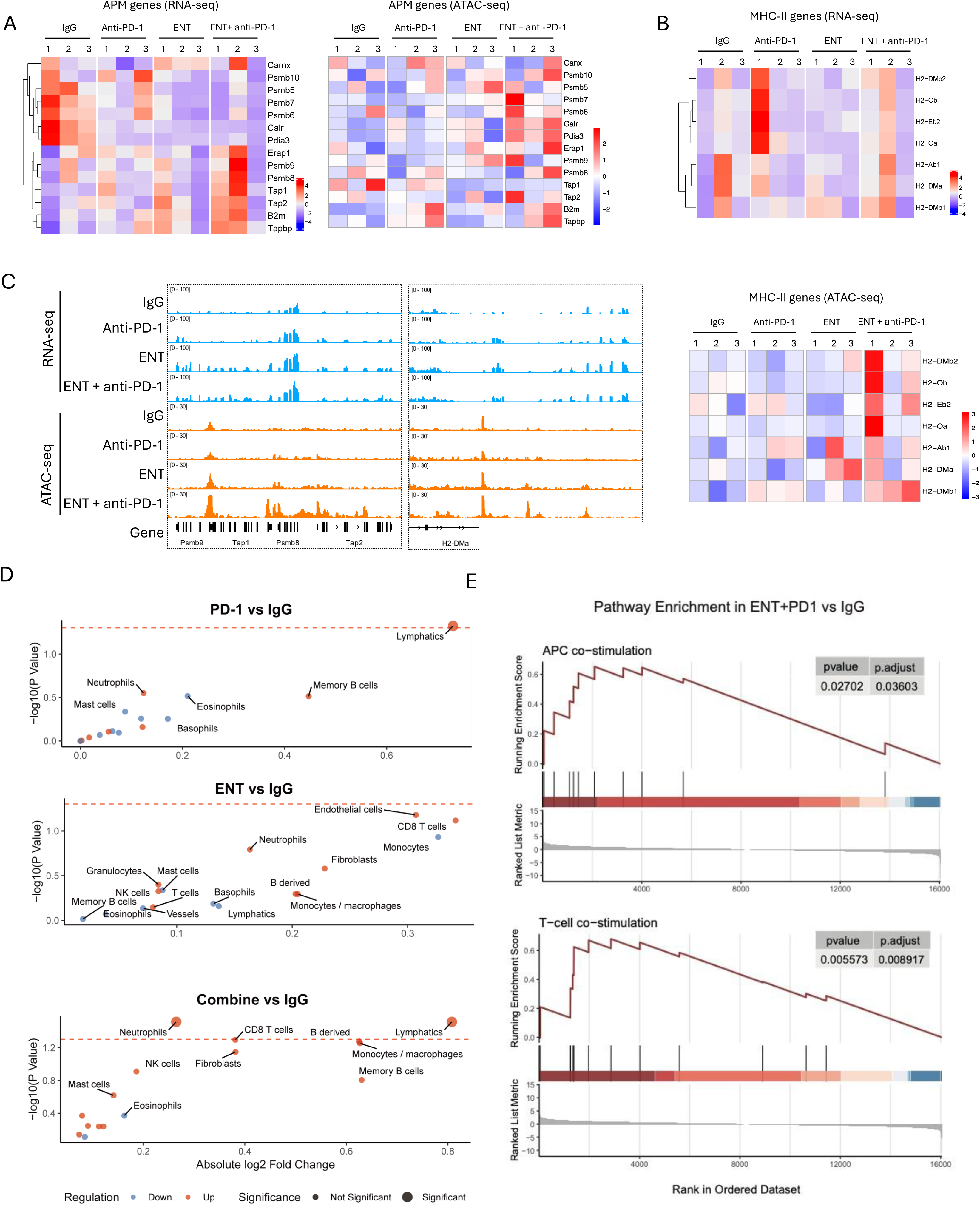
Entinostast regulates APM genes and MHC-II genes, enhancing T-cell immune response A. Heatmap of antigen presentation machinery genes on their expression level and ATAC-seq peak signal in these gene’s promoter region (2kb upstream and downstream of TSS) B. Heatmap of MHC-II genes on their expression level and ATAC-seq peak signal in these gene’s promoter region (2kb upstream and downstream of TSS) C. RNA-seq peaks and ATAC-seq peaks track signals on the APM genes and MHC-II genes. D. Comparison between each treatment groups and IgG for the immune cell infiltration differences with limma method. E. Gene Set Enrichment Analysis (GSEA) between ENT + anti-PD-1 and IgG group.

Activation of APM genes is associated with the restoration of MHC-I expression on the tumor cell surface for CD8^+^ T cell recognition.^42,43^ Effective antigen presentation and immune activation also require the participation of antigen-presenting cells (APCs) during an adaptive immune response. MHC-II molecules are predominantly expressed in APCs, including dendritic cells, macrophages, and B cells. We examined the chromatin accessibility of MHC-II genes in tumors using ATAC sequencing and observed MHC-II gene activation and chromatin opening at MHC-II gene loci with treatment (Fig. 5B-C and Fig. S5A), suggesting priming of immune cells to recognize tumors.

### Entinostat plus anti-PD-1 primes the tumor immune microenvironment

To further evaluate the impact of entinostat and anti-PD1 treatments on the tumor microenvironment, we performed immune deconvolution analysis using CIBERSORT ^44,45^ and observed different immune cell composition across the treatment groups (Fig. S5B). We next used limma method^46^ to compare the differences in immune cell infiltration between the treatments and found that both entinostat alone and in combination with anti-PD1 upregulated CD8^+^ T cell infiltration. The PD-1 monotherapy did not increase CD8+ T cell infiltration (Fig. 5D). Given that the entinostat plus anti-PD-1 group showed the most pronounced changes, we performed Gene Set Enrichment Analysis (GSEA) between entinostat alone and the combination and observed significant enrichment of multiple immune-related pathways, including “T Cell Co-Stimulation” (NES = 1.82, adjusted p-value = 0.01) and “Apc Co-Stimulation” (NES = 1.61, adjusted p-value = 0.04) upon the combination treatment, suggesting that entinostat plus anti-PD-1 synergistically promotes anti-tumor immune responses by enhancing T cell activation and improving antigen presentation efficiency (Fig. 5E).

### Entinostat plus anti-PD-1 enhances T-cell infiltration and cytotoxicity in tumors

To assess T-cell infiltration upon entinostat and anti-PD-1 treatments, we carried out immunostaining for T-cell markers using the necropsy samples. We found that the CD3^+^ and CD8^+^ immune infiltrates in vehicle-treated control tumors are localized peritumorally and in proximity to vascular structure or airway in the tumor (Fig. 6A). Entinostat plus anti-PD-1 led to an increase in intratumoral CD3^+^, CD4^+^, and CD8^+^ T cells by IHC and quantitatively (Fig. 6A-B). Moreover, pro-inflammatory cytokines such as *CXCL11, CXCR3* and *IL-6* were also elevated upon entinostat and the combination treatments (Fig. 6C), indicating enhanced chemokine-mediated T-cell recruitment.

**Figure 6.**
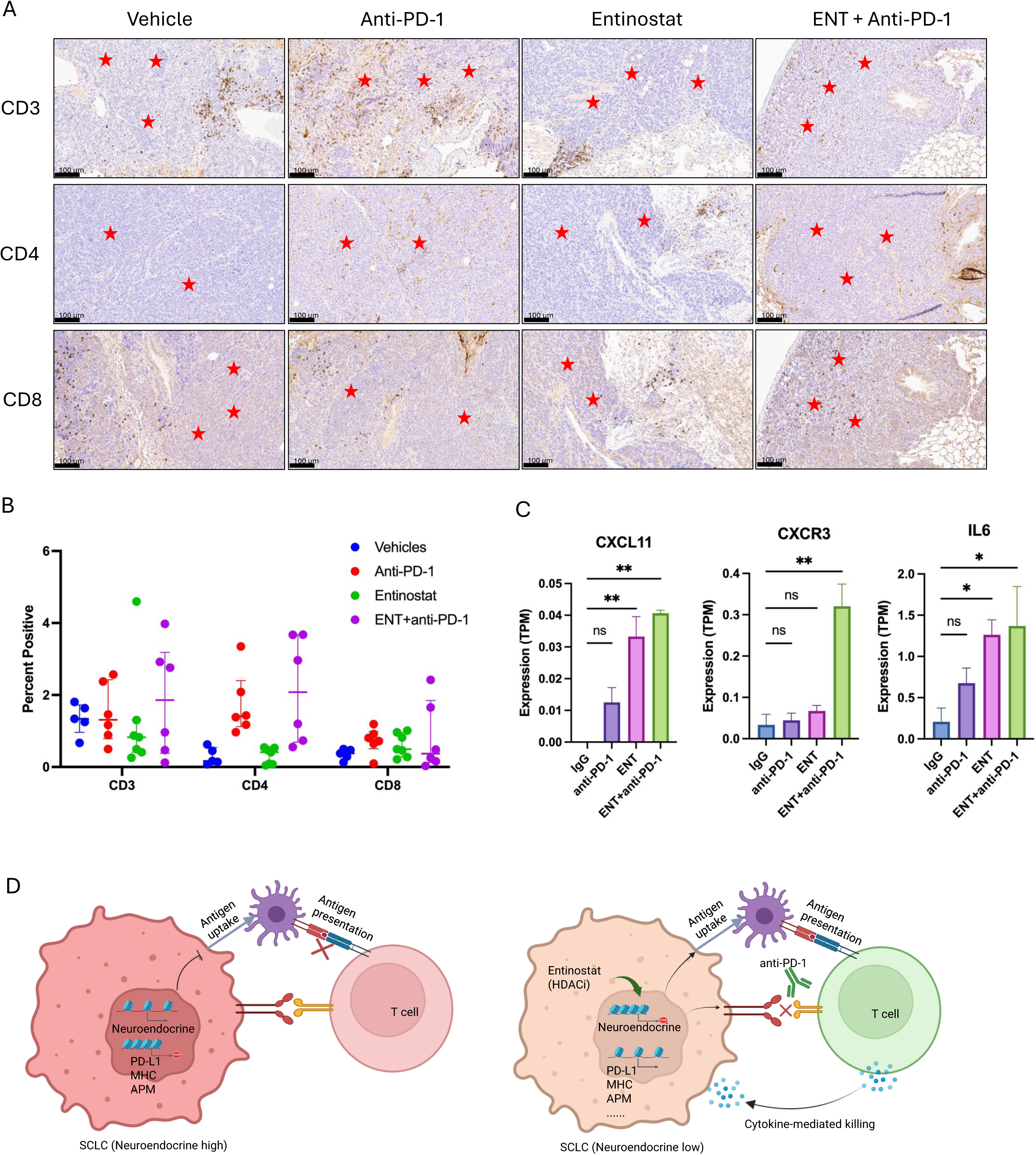
Entinostat plus anti-PD-1 enhances T-cell infiltration and cytotoxicity in tumors. A. IHC staining of immune cell markers *CD3, CD4* and *CD8*. Scale bar is uM (20x). red star marks tumor regions. B. Quantification of IHC staining on immune cell markers *CD3, CD4, CD8* positive cells. C. Gene expression for the chemokine genes. TPM: Transcripts Per Million. D. Model for entinostat function in SCLC immune therapy. Left: SCLC in neuroendocrine-high status have NE-high associated genes highly expressed while MHC genes and APM genes repressed. Antigen processing is blocked. Anti-PD-1 response is limited. T-cell infiltration was minor and not active to kill tumor cell. right: entinostat can regulate key neuroendocrine genes and change NE-high to NE-low while activate MHC and APM genes, enable antigen processing and work together with anti-PD-1 to facilitate the activity of T-cells in killing tumor cells.

In summary, Myc-driven RPM SCLC exhibits NE high phenotype and low immunogenicity, marked by minimal antigen presentation and low PD-L1 expression (Fig. 6D, left). HDAC inhibition unlocks tumor plasticity by reducing NE characteristics and epigenetically activating APM genes. By altering the tumor microenvironment, HDAC inhibition leads to enhanced antigen presentation and enables effective anti–anti-PD-1-mediated tumor clearance through enhanced T cell-mediated infiltration and cytotoxic activity (Fig. 6D, right).

### Entinostat plus anti-PD-1 improves survival in the RPM allografts

The combination of entinostat and anti-PD-1 treatment did not lead to a significant reduction in tumor volume and increase in survival in the primary RPM model. The proximity of primary lung tumors to vital structures, such as the bronchus, which necessitated early euthanasia and may have limited the time window for immunotherapy. Therefore, we established an allograft model in which RPM tumors were transplanted subcutaneously into immune-competent, strain-matched recipients. These tumors were allowed to grow for 3 to 5 weeks before undergoing treatment with entinostat, anti-PD-1, or the combination (Fig. 7A). Combination treatment with daily oral entinostat and bi-weekly IP anti-PD-1 increased tumor growth inhibition over either treatment alone, and significantly improved survival (Fig 7B – C) compared to vehicle, isotype control, and anti-PD1 (p = 0.0257, 0.0015, and 0.0058, respectively), validating the therapeutic benefit of this approach in a more tractable system.

**Figure 7.**
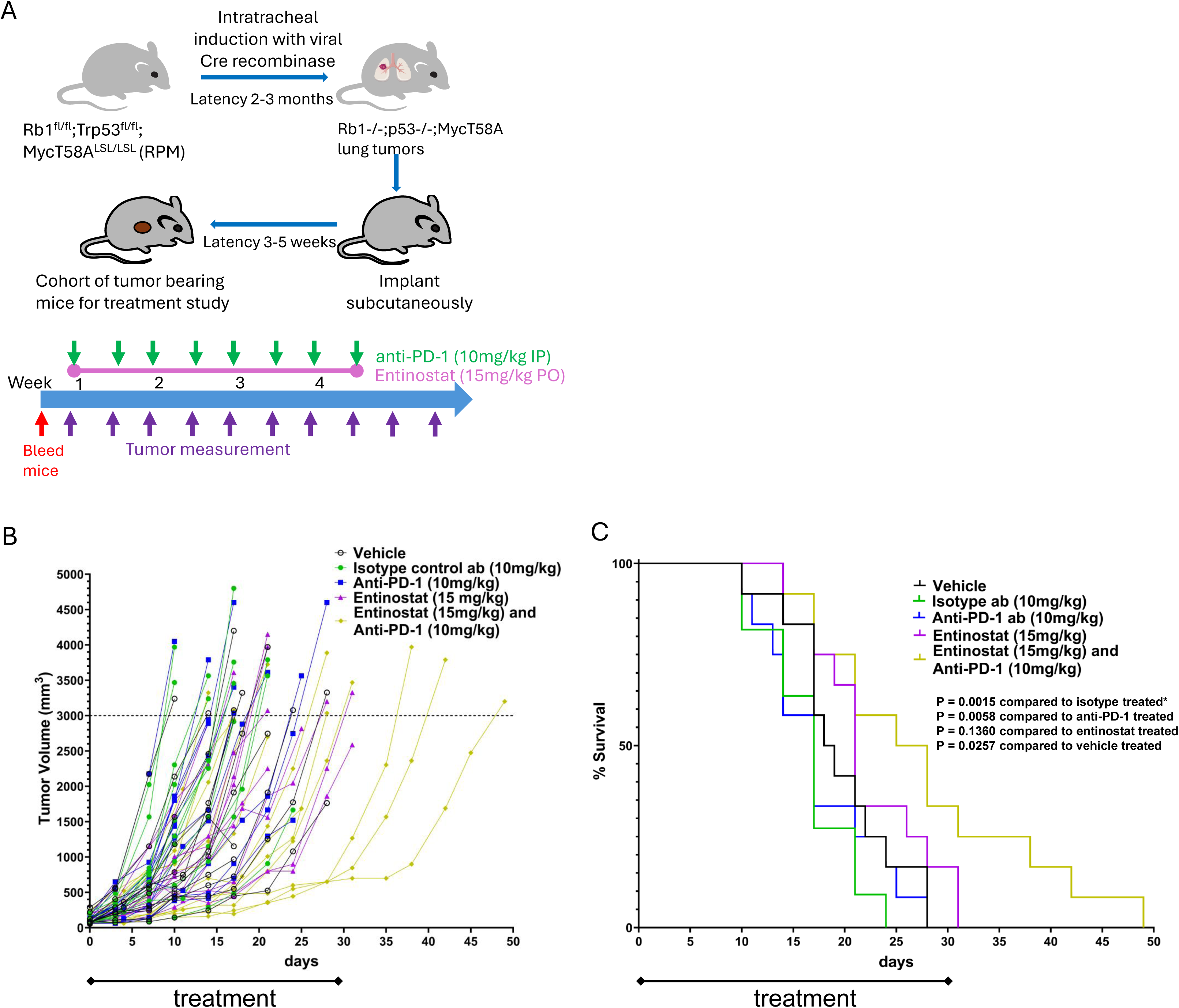
Entinostat plus anti-PD-1 improves survival in the RPM allografts A. Scheme of SCLC RPM allograft mouse model. The GEM *Rb1/Trp53/Myc^T58A^* (RPM) mice was intratracheal injected with viral Cre recombinase and latency after 2 - 3 months when it generates lung tumors. The tumors were transplanted subcutaneously to strain-matched and immunecompetent recipients. After 3 – 5 weeks, treatment commenced according to the study timeline. B. Tumor volume record of allograft mouse control and treatment groups. C. Survival curve of allograft mouse control and treatment groups.

## DISCUSSION

SCLC is characterized by aggressive growth, early metastasis, and frequent relapse following standard chemo-immunotherapy. Its molecular subtypes differ in therapeutic vulnerability—NE-high tumors respond better to chemotherapy, while NE-low subtypes demonstrate increased immune infiltration and are more responsive to immune checkpoint blockade.^47^ Hence, tailoring therapies based on subtype-specific (NE status and immune status) could be developed as a strategy to improve outcomes.

Here, we demonstrate that the class I HDAC inhibitor entinostat epigenetically reprograms NE-high, immune-cold SCLC toward an NE-low, immune-responsive state. First, we show that in human SCLC cell lines, entinostat induces a robust, dose-dependent upregulation of PD-L1, MHC molecules, and antigen processing machinery gene expression-key features required to elicit an effective antitumor immune response. Next, using immunocompetent RPM genetically engineering mouse models (GEMMs), we discovered that primary SCLC tumors suppress immune (PDL-1 and MHC) and APM gene expression relative to normal lung tissue, suggesting a mechanism of immune evasion. Notably, treatment with entinostat, either alone or in combination with anti-PD-1 therapy, reversed this suppression and significantly upregulated immune gene expression, enhancing tumor immunogenicity.

In addition to immune reprogramming, entinostat induced a phenotypic shift of MYC-driven RPM tumors from NE-high to NE-low, characterized by downregulation of canonical NE markers (e.g., *Neurod1*, *Ascl1*, *Syp*) and upregulation of NE-low lineage markers (*Yap1*, *Pou2f3*). ATAC-seq analysis revealed that entinostat increased chromatin accessibility at immune-related and lineage-defining loci, likely through indirect effects on acetylated transcription factors.^48^ Finally, the combination of entinostat and anti-PD-1 therapy yielded synergistic effects on increased T-cell infiltration, suppressed tumor growth, and improved survival in RPM allograft models. These findings are consistent with prior reports demonstrating that HDAC inhibition enhances the efficacy of immune checkpoint blockade by mitigating suppressive myeloid populations and promoting T-cell–mediated antitumor immunity.^20^

While prior clinical trials (e.g., ENCORE 601 in non-small cell lung cancer, NSCLC) showed limited benefit of entinostat–pembrolizumab combination, the combination did provide a clinical benefit with higher circulating classical monocytes.^49^ SCLC is inherently driven by epigenetic regulation and exhibits greater lineage plasticity compared to NSCLC, making it particularly susceptible to epigenetic modulation as a means to unlock therapeutic potential.^50,51^

In conclusion, these findings underscore the potential of entinostat as a potent epigenetic modulator that reprograms tumor cell-intrinsic properties in SCLC by inducing a durable phenotypic shift from NE-high to NE-low, while simultaneously enhancing tumor immunogenicity through restoration of antigen presentation and promotion of immune cell infiltration.This dual effect on tumor-intrinsic and immune-modulating pathways highlights entinostat as a promising component of combination immunotherapy regimens in SCLC. Clinical trials are warranted to assess their translational potential in this setting. Future studies should also focus on elucidating the detailed molecular mechanisms by which entinostat regulates the neuroendocrine differentiation and immune modulation (i.e., the activation and recruitment of transcription factors that regulate PD-L1 and MHC expression induced by HDAC inhibition).

## Resource availability

### Lead contact

Further information and requests for resources and reagents should be directed to and will be fulfilled by the lead contact, Azam Ghafoor.

## Acknowledgments

This work was in part supported by NIH grants 4R00CA273171-02 (to Y.S.) and ZIA BC 011793 (to A.T). This research was supported by the Intramural Research Program of the National Institutes of Health (NIH). The contributions of the NIH author(s) were made as part of their official duties as NIH federal employees, are in compliance with agency policy requirements, and are considered Works of the United States Government. However, the findings and conclusions presented in this paper are those of the author(s) and do not necessarily reflect the views of the NIH or the U.S. Department of Health and Human Services. This project has been funded in part with Federal funds from the National Cancer Institute, National Institute of Health, under Contract No. HHSN26120150003I. The content of this publication does not necessarily reflect the views or policies of the Department of Health and Human Services, nor does mention of trade names, commercial products, or organizations imply endorsement by the U.S. Government

## Materials and Methods

### Small cell lung cancer cell lines

H82 cancer cell was provided by Dr. Haobin Chen’s lab (NCI, Bethesda, MD) and H524, H889, H209, and DMS-114 cell lines were purchased from American Type Culture Collection (Manassas, VA). Cells were maintained in culture in RPMI-1640 medium supplemented with 10% FBS, penicillin/streptomycin 1x, and glutamax 1x. Human cell lines H82 and H524 are variant forms of human SCLC (high-myc expression) whereas H524 and H209 are classic forms (low-myc expression).

### Reagents

Entinostat (99% purity) used for animal studies was purchased from MedChemExpress LLC and formulated in in 0.5% methycellulose solution in saline. Instructions included measuring and placing an appropriate amount of entinostat at the bottom of a tube (15ml or 50ml), adding 0.5% of methycellulose on top, and then sonicating to disperse entinostat in the 0.5% methycellulose into a suspension. Methylcellulose (400) M0262-100G was purchased from Sigma Aldrich.

Entinostat was made fresh weekly, and it is stored at 4°C between uses and kept on ice during usage. Entinostat for in vitro use was prepared in DMSO at 10mM stock. Treatment antibodies for in vivo use were anti-PD-1 antibody obtained from Bio X Cell (RPM1-14, rat IgG2a) and isotype control antibody (2A3, Rat IgG2a) obtained from Bio X Cell. For IHC analysis, CD3 (Bio-Rad cat. #MCA1477), CD4 (Thermo-Fisher cat. #13-9766-82), CD8 (Thermo-Fisher cat. #14-0195-82), PD-L1 (R & D Systems cat. #AF1019), and MHCI (Bio-Rad cat. #MCA2397), MHC II (Abcam cat. #ab25333), and CD56 (Sigma cat. #HPA039835).

### RNA isolation, reverse transcription, and real-time PCR

Cells were pelleted and RNA were isolated using RNeasy Mini kit (QIAGEN, Germantown, MD) according to the manufacturer’s protocol. RNA was quantified using NanoDrop 2000 (Thermo-Fisher, Waltham, MA) and frozen at-70°C before undergoing reverse transcription. RNA was reverse transcribed using the High-Capacity cDNA Reverse Transcription Kit (Life Technologies, CA) in an Applied Biosystems GeneAmp PCR system 9700 (Life Technologies).Quantitative real-time PCR (qPCR) was carried out in a StepOnePlus real time PCR instruments (Life Technologies) using a 96-well reaction plate and TaqMan Fast Advanced Master Mix as well as primers and probe for each assay (Life Technologies). TaqMan real time PCR assays (HLA-A, HLA-B, HLA-C, PD-L1, HPRT, and PGK1 were purchased from Life Technologies. Delta Ct value was calculated as Ct value of target gene probe subtracted by the geometric mean of Ct values of HPRT and PGK1 probes. The relative gene expression level was calculated as 2-deltaCt using assays for HPRT and PGK1 as the internal controls.

### Western Blotting

2 × 10^6^ SCLC cells were seeded and treated with Entinostat of indicated concentrations for 24 hours or with 0.5 μM Entinostat for indicated time points. Cells were collected and lysed with RIPA buffer (150 mM NaCl, 1% Nonidet P-40, 0.5% sodium deoxycholate, 0.1% SDS and 25 mM Tris pH 7.4, 1 × protease inhibitor cocktail (Cell Signaling, #5871), 1 mM DTT), followed by sonication (40% output for 10 sec pulse and 10 sec rest for 4 times. Cellular proteins were quantitated by Bradford protein assay using SoftMax Pro 7.1. 10 μg whole cellular lysates were resolved by SDS-PAGE electrophoresis using 4–20% Mini-PROTEAN® TGX™ precast gels (Bio-Rad, #4561096), followed by transfer to PVDF membranes using Trans-Blot® Turbo™ system (Bio-Rad, #1704150). Blots were blocked 1X TBS-T with 5% w/v nonfat dry milk for 1 h then incubated with primary antibodies in 1X TBS-T with 5% w/v nonfat dry milk at 1:1,000 dilution overnight at 4 °C. Primary antibodies included anti-mouse PD-L1 (405.9A11) monoclonal antibody (1:500, Cell Signaling Technology), anti-rabbit PD-L1 (E1L3N) monoclonal antibody (1:500, Cell Signaling Technology), anti-acetylated H-3 antibody (positive control, 1:1000, CST), and β-actin (loading control). After 1h incubation with secondary antibodies 1X TBS-T with 5% w/v nonfat dry milk at 1:10,000 dilution for 1 h at room temperature, blots were developed using SuperSignal™ West Femto ECL substrates (Thermo Fisher, #34096) and signals were detected by ChemiDoc^TM^ MP Imaging System (Bio-Rad). anti-mouse PD-L1 (405.9A11) monoclonal antibody (1:500, Cell Signaling Technology), anti-rabbit PD-L1 (E1L3N) monoclonal antibody (1:500, Cell Signaling Technology), anti-acetylated H-3 antibody (positive control, 1:1000, CST), and β-actin (loading control). Quantitative measurements were performed using GraphPad (Prism 7) software. Western blot analysis was conducted were used in at least two independent experiments

### Animals

All animal studies were approved by the National Institutes of Health Institutional Animal Care and Use Committee. Breeder pairs of the Rb1 ^fl/fl^;p53 ^fl/fl^;Myc ^lsl/lsl^ T58A SCLC GEM (Igs2tm1(CAG-Myc*T58A/luc)Wrey Trp53tm1Brn Rb1tm3Tyj/OlvrJ; stock no. 029971) were obtained from The Jackson Laboratory (Bar Harbor, ME). For initial evaluation, SCLC was induced in a cohort of 20 mice at 8 weeks of age by intratracheal instillation of 2.5 × 107 PFU of Ad5CGRP-Cre (Product #VVC-Berns-1160, U. of Iowa Viral Vector Core Facility). Mice were weighed weekly and observed daily during the study to monitor for signs of tumor development. For the treatment efficacy study, a total of 66 RPM mice were produced in 3 cohorts and induced as described above. Tumor bearing mice (confirmed by MRI) were randomized by volume into four experimental groups: vehicle (control), entinostat monotherapy (15 mg/kg daily for 28 days by oral gavage (PO)), anti–PD-1 antibody monotherapy (10 mg/kg of anti-PD-1 delivered by intraperitoneal injection (IP) twice weekly for four weeks), and combined entinostat and anti– PD-1 antibody treatment. 0.5% methylcellulose (entinostat vehicle) and isotype control (Rat IgG2a) antibody treatment groups served as controls.

For the efficacy study in an RPM allograft model, For the RPM allograft model, tumors were harvested from induced RPM mice and tumor fragments were re-implanted subcutaneously into uninduced RPM mice. For the treatment study, passage 3 subcutaneous tumor was dissociated using manual dissection and incubation in a mixture of DMEM, trypsin and collagenase. The freshly dissociated cells were injected subcutaneously into uninduced RPM GEM mice (5 × 104 cells in 0.2ml per mouse) to create the treatment cohort. The flank was palpated twice weekly in order to detect tumor growth at the earliest stages. Mice were recruited to the study as tumors became palpable and measurable by caliper and distributed into treatment groups such that the average volume was ∼100mm3 per group. Mice were euthanized when tumor volume reached or exceeded 3000mm3 (L xW2/2) or if any clinical endpoint signs were noted.

### MRI

Magnetic resonance imaging (MRI) was used to calculate 3D tumor volumes. Mouse lungs were imaged in the supine position on an MRI 3.0T clinical scanner (Philips Healthcare Intera Achieva). Lung tumor volumes were measured in mm3 using ITK-SNAP software (Version 2.2.0) for image analysis.

### Necropsy

Mice were euthanized according to humane endpoints in a protocol approved by NCI’s Animal Care and Use Committee. Indications included body weight loss and clinical signs of advanced tumors such as slowed breathing. For initial characterization, the entire lung was harvested and inflated with formalin to evaluate location and context of tumor nodules within the lung For evaluation of tumors in the drug treatment study, tumors were harvested at necropsy and divided into sections fixed in formalin for histopathology analysis or flash-frozen for RNA-seq and ATAC-seq.

### Histopathology analysis

Hematoxylin and eosin (H&E) staining was performed on fixed paraffin-embedded tissue and analyzed by a board-certified DVM pathologist (L.B.). Select tumors were stained for CD56 (NCAM; Sigma cat. #HPA039835), PD-L1 (R & D Systems cat. #AF1019), MHC I (Bio-Rad cat. #MCA2397), MHC II (Abcam cat. #ab25333), CD3 (Bio-Rad cat. #MCA1477), CD8

(Thermo-Fisher cat. #14-0195-82), and CD4 (Thermo-Fisher cat. #13-9766-82). All immunohistochemical stains were performed on fixed paraffin embedded tissue. Slides were scanned at 20X on an Aperio (Leica) microscope scanning/analysis system. For each slide a semi-automated quantification was employed using the Aperio image analysis software.

### RNA purification for RNA-seq analysis

Flash-frozen tumor tissue was manually dissected with a scalpel blade and then fully homogenized using a TissueLyzer. After centrifugation, supernatant from the tissue lysates was transferred to an AllPrep DNA spin column (Qiagen) for DNA extraction. Flow-through from the column was transferred to a RNeasy spin column (Qiagen) for RNA purification. DNA was then eluted from the DNA spin columns and reserved for later use. For ATAC-seq, flash-frozen tumors were sent to the sequencing facility for DNA purification and sequencing.

### Bioinformatics data analysis

For RNA-seq, the raw FASTQ files were trimmed using timGalore (version 0.4.5, https://www.bioinformatics.babraham.ac.uk/projects/trim_galore/) and Trimmomatic (version 0.36).(Bolger et al., 2014) Surviving reads were aligned to the mm9 genome with the STAR aligner version 2.5.4a).^52^ using the UCSC mm9 gene annotation using the two-pass mode and “best recall at base and read level” settings as shown in Table S37 from.^53^ Gene expression raw read counts were quantified using TPMcalculator (version 0.0.3)^54^ “ExonReads” column.

Differential expression analysis was performed using DESeq2 (version 1.22.2)^55^ in R (version 3.5.1).

For ATAC-seq analysis, the raw FASTQ files were trimmed using timGalore, aligned to the mm9 genome with the BWA.^56^ Deeptools was used to generate the bigwig files to view in IGV. Peaks were called using MACS. Heatmaps were generated in R with pheatmap function.

NE score calculation was followed the published method.^40^ In this method, a 50-gene expression signature is used. This signature consists of 25 genes known to be upregulated in neuroendocrine (NE) cells and 25 genes upregulated in non-NE cells. The NE score quantifies the degree to which a cell line expresses the NE-specific genes relative to the non-NE genes. A quantitative NE score was generated from this signature using the formula: NE score = (correlation NE – correlation non-NE)/2 where correlation NE (or non-NE) is the Pearson correlation between expression of the 50 genes in the test sample and expression of these genes in the NE (or non-NE) cell line group.

For immune infiltration analysis, we used mMCPcounter (version 1.1.0)^39^ to evaluate the immune cell infiltration in the RPM model. To validate the mMCP results, we also applied CIBERSORT (version 0.1.0)^44^ with a list of mouse immune cell signatures.^45^ The limma method^46^ was used to compare the differences in immune cell infiltration between three treatment groups and the control group. All results were visualized using ggplot2 (version 3.5.2) [Wickham H (2016). ggplot2: Elegant Graphics for Data Analysis. Springer-Verlag New York. ISBN 978-3-319-24277-4, https://ggplot2.tidyverse.org.] and ggpubr (version 0.6.0) [Kassambara, Alboukadel. “ggpubr:’ggplot2’ based publication ready plots.” R package version (2018): 2.]. For Gene Set Enrichment Analysis (GSEA), we performed homology conversion of the Bulk RNA-seq data from RPM using the biomaRt (version 2.58.2).^57^ Then we performed pathway enrichment analysis using GSVA(version 2.0.7)^58^ with public immune gene sets.^59^

### Statistics

Statistical analyses were performed in GraphPad Prism 7 (GraphPad Software) and listed in figure legends. Statistical significance was set at p<0.05. *p<0.05, **p<0.01, ***p<0.001, ****p<0.0001.

### Study approval

All animal studies were conducted under approval of the NCI Animal Care and Use Committee. NCI-Frederick is accredited by AAALAC International and follows the Public Health Service Policy for the Care and Use of Laboratory Animals. Animal care was provided in accordance with the procedures outlined in the ‘Guide for Care and Use of Laboratory Animals (National Research Council; 2011; National Academy Press; Washington, D.C.).

