## Supplemental Figures for "HDAC inhibition unlocks tumor plasticity and enhances immunotherapy response in Myc-Driven Small Cell Lung Cancer"

**Figure legend**

**Supplementary Figure**

Figure S1. Immune checkpoint and MHC genes expression and it's correlation with DNA methylation.

A. Gene expression levels of MHC-I genes including *HLA-A, HLA-B, HLA-C* and *TAP1* in SCLC and lung adenocarcinoma. Data was from sclcCellMinerCDB. **** p < 0.0001.
B. Correlation between promoter DNA methylation signal and gene expression of *PD-L1 (CD274), HLA-B* in SCLC cell lines. Plot is generated from the SclcCellMinerCDB.

Figure S2. Gene expression of MHC-II, chemokine genes and antigen presenting machinery genes in SCLC cell lines.

A. The qRT-PCR for gene expression of MHC-II genes (*HLA-DRA, HLA-DQ1 HLA-DP*) and MHC-I gene (*MICA*) on SCLC cell lines H889, H209, H82, H524, and DMS-114 treated with Entinostat for 24 hours in increasing dose of 0uM, 0.2uM, 1uM and 5uM. Relative gene expression values are normalized to two control gene HPRT and PGK1 and shows at a 100 fold manner. Error bars are representative of technical triplicates. *p<0.05, **p<0.01, ***p<0.001, ****p<0.0001.
B. The qRT-PCR for gene expression of chemokine genes (*CXCL10, IFNγ*) same as treatment in A.
C. The qRT-PCR for gene expression of antigen presenting machinery genes (TAP1, PSMB8) same as treatment in A.

Figure S3. RPM mouse model grwoth and HE staining.

A. MRI examples for the RPM mouse model to show the tumor growth.
B. HE stains of RPM mouse tumors to show the features of SCLC: small size cells, scant cytoplasm, no distinct nucleoli, and neoplastic cells arranged in sheets and ribbons. Left panel: tumor arise in the large bronchi. Right top panel: 5X view of tumor adjacent to the large airway. Right bottom panel: 20X view of SCLC cells.
C. HE stains of RPM tumors to show lobular coagulative necrosis consistent with tumor-associated vascular occlusion (arrow). Occlusion and necrosis were observed in some cases even when tumors were not widespread.

Figure S4. Characterize the toxicity and tumor growth of entinostat on RPM tumors.

A. Western blot to show the kinetics of PD-L1 expression in H889 cell line during 24-hour treatment with 0.5 uM entinostat and following removal from the culture medium for 24 hours.
B. Tolerance test of entinostat and in combination use with anti-PD-1 by mouse body weight measurement. Entinostat treatment occurred on day 1-7 and anti-PD-1 treatment was given on day 1 and day 4.
C. Tumor volume record for each sample by MRI.

D. MRI examples for each group at initial and late stages.

Figure S5. Entinostat effect on APM genes and immune cells.

A. RNA-seq peaks and ATAC-seq peaks track signals on the APM genes.
B. Immune cell infiltration analysis on the four sample groups with CIBERSORT.

Fig S1

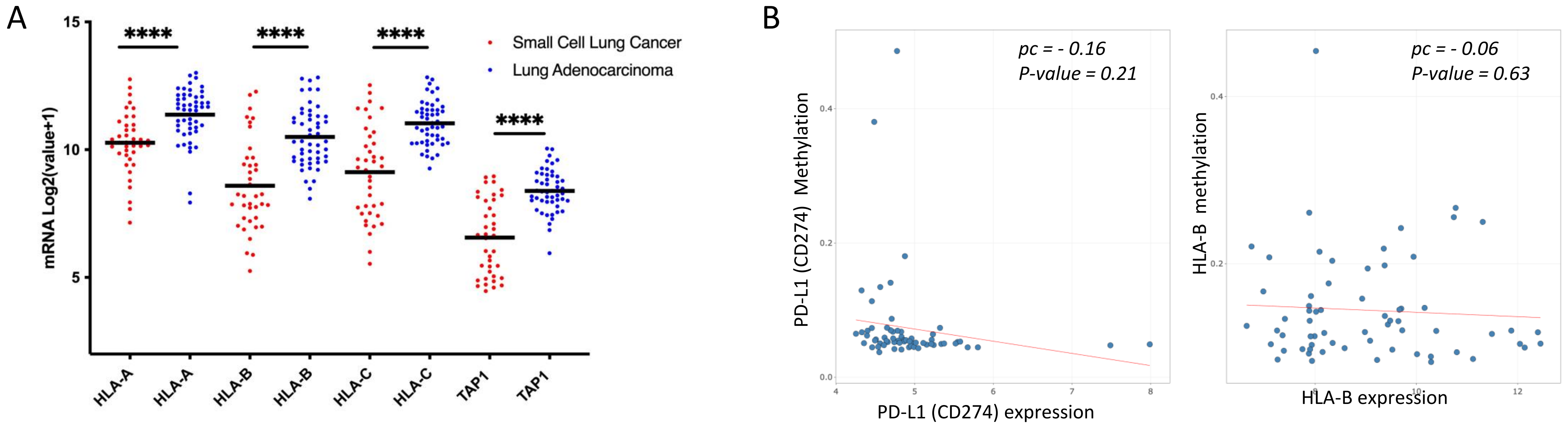
A
15
10
5
mRNA Log2(value+1)
****
****
****
****
Small Cell Lung Cancer
Lung Adenocarcinoma
HLA-A
HLA-A
HLA-B
HLA-B
HLA-C
HLA-C
TAP1
TAP1
B
pc = - 0.16
P-value = 0.21
PD-L1 (CD274) Methylation
PD-L1 (CD274) expression
pc = - 0.06
P-value = 0.63
HLA-B methylation
HLA-B expression

Fig S2

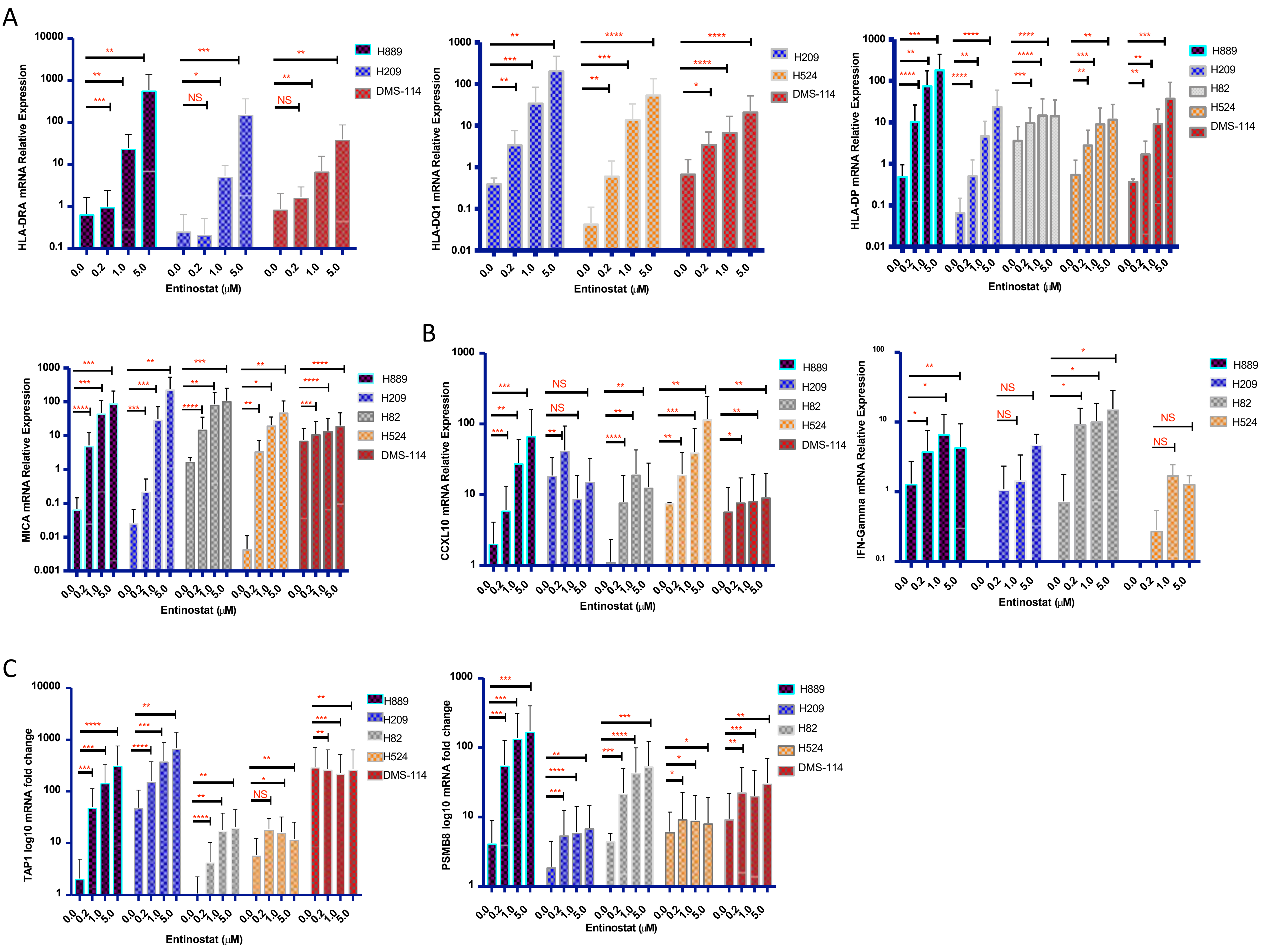
A
HLA-DRA mRNA Relative Expression
HLA-DQ1 mRNA Relative Expression
HLA-DP mRNA Relative Expression
MICA mRNA Relative Expression
B
CCXL10 mRNA Relative Expression
IFN-Gamma mRNA Relative Expression
C
TAP1 log10 mRNA fold change
PSMB8 log10 mRNA fold change
Entinostat (μM)
0.0
0.2
1.0
5.0
H889
H209
H82
H524
DMS-114
NS

Fig S3

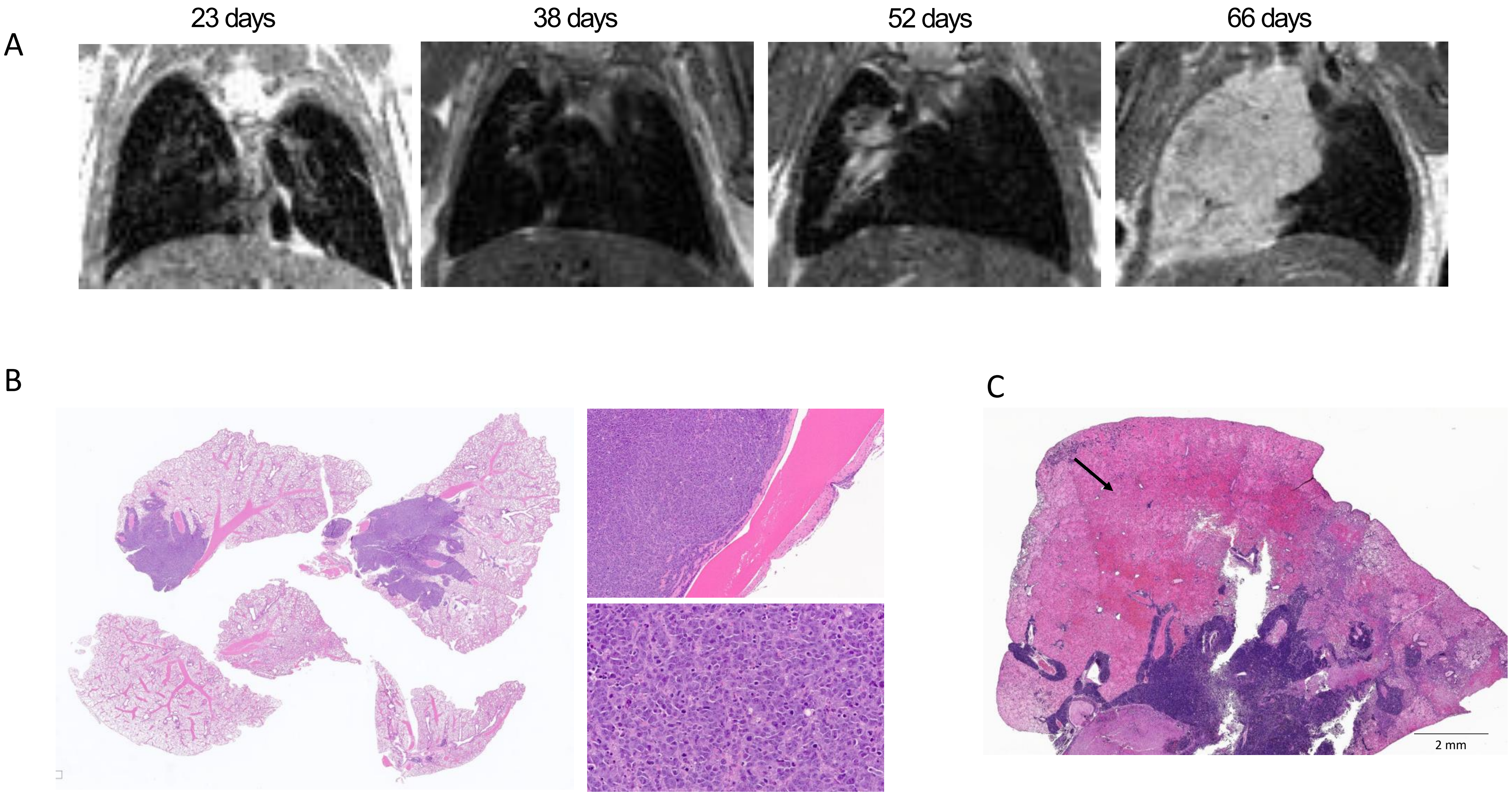

Fig S4

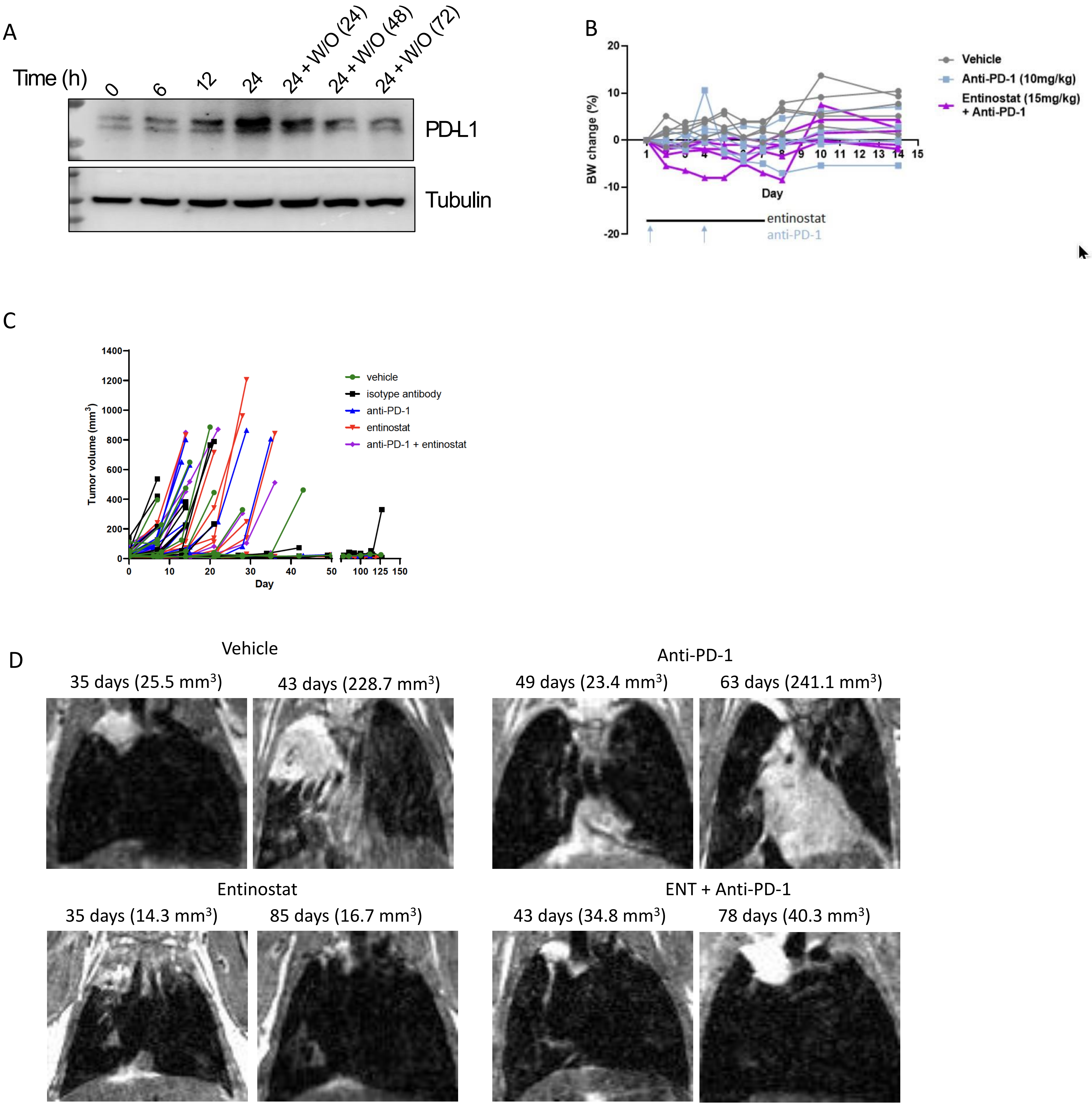

A
Time (h)
0
6
12
24
24 + W/O (24)
24 + W/O (48)
24 + W/O (72)
PD-L1
Tubulin
B
BW change (%)
Day
Vehicle
Anti-PD-1 (10mg/kg)
Entinostat (15mg/kg) + Anti-PD-1
entinostat
anti-PD-1
C
Tumor volume (mm3)
Day
vehicle
isotype antibody
anti-PD-1
entinostat
anti-PD-1 + entinostat
D
Vehicle
35 days (25.5 mm3)
43 days (228.7 mm3)
Anti-PD-1
49 days (23.4 mm3)
63 days (241.1 mm3)
Entinostat
35 days (14.3 mm3)
85 days (16.7 mm3)
ENT + Anti-PD-1
43 days (34.8 mm3)
78 days (40.3 mm3)

Fig S5

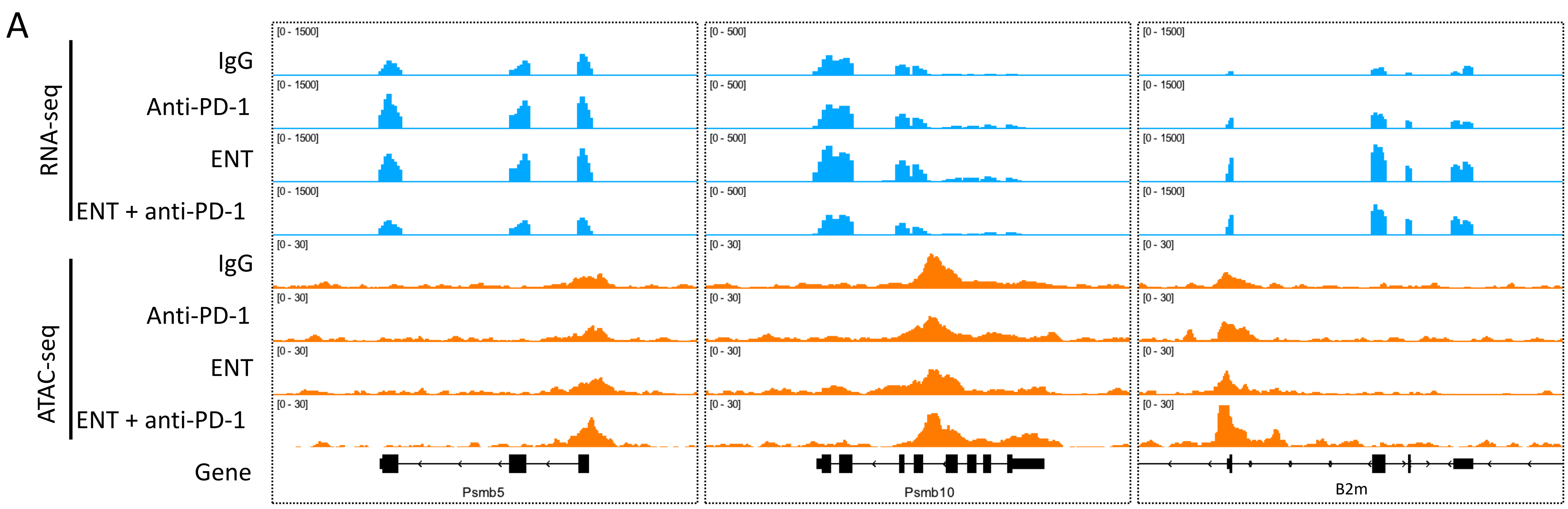
A
RNA-seq
ATAC-seq
IgG
Anti-PD-1
ENT
ENT + anti-PD-1
IgG
Anti-PD-1
ENT
ENT + anti-PD-1
Gene
[0 - 1500]
[0 - 500]
[0 - 30]
Psmb5
Psmb10
B2m

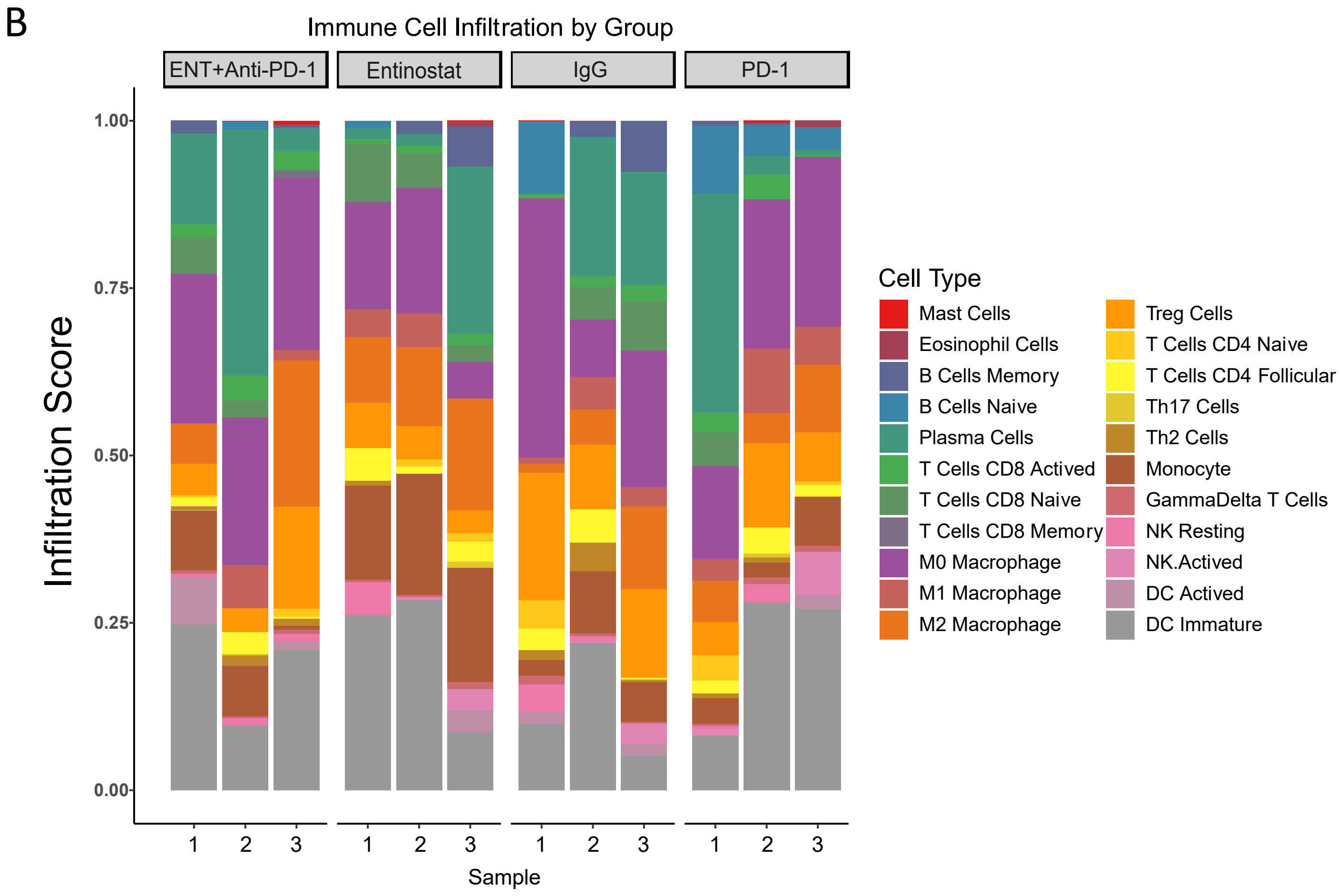
B
Immune Cell Infiltration by Group
ENT+Anti-PD-1
Entinostat
IgG
PD-1
Infiltration Score
1.00
0.75
0.50
0.25
0.00
1 2 3
Sample
Cell Type
Mast Cells
Eosinophil Cells
B Cells Memory
B Cells Naive
Plasma Cells
T Cells CD8 Actived
T Cells CD8 Naive
T Cells CD8 Memory
M0 Macrophage
M1 Macrophage
M2 Macrophage
Treg Cells
T Cells CD4 Naive
T Cells CD4 Follicular
Th17 Cells
Th2 Cells
Monocyte
GammaDelta T Cells
NK Resting
NK.Actived
DC Actived
DC Immature
